# Poly-Enrich: Count-based Methods for Gene Set Enrichment Testing with Genomic Regions

**DOI:** 10.1101/488734

**Authors:** Christopher T Lee, Raymond G Cavalcante, Chee Lee, Tingting Qin, Snehal Patil, Shuze Wang, Zing TY Tsai, Alan P Boyle, Maureen A Sartor

## Abstract

Gene set enrichment (GSE) testing enhances the biological interpretation of ChIP-seq data and other large sets of genomic regions. Our group has previously introduced two GSE methods for genomic regions: ChIP-Enrich for narrow regions and Broad-Enrich for broad genomic regions, such as histone modifications. Here, we introduce new methods and extensions that more appropriately analyze sets of genomic regions with vastly different properties. First, we introduce Poly-Enrich, which models the number of peaks assigned to a gene using a generalized additive model with a negative binomial family to determine gene set enrichment, while adjusting for gene locus length (#bps associated with each gene). This is the first method that controls for locus length while accounting for the number of peaks per gene and variability among genes. We also introduce a flexible weighting approach to incorporate region scores, a hybrid enrichment approach, and support for new gene set databases and reference genomes/species.

As opposed to ChIP-Enrich, Poly-Enrich works well even when nearly all genes have a peak. To illustrate this, we used Poly-Enrich to characterize the pathways and types of genic regions (introns, promoters, etc) enriched with different families of repetitive elements. By comparing ChIP-Enrich and Poly-Enrich results from ENCODE ChIP-seq data, we found that the optimal test depends more on the pathway being regulated than on the transcription factor or other properties of the dataset. Using known transcription factor functions, we discovered clusters of related biological processes consistently better modeled with either the binary score method (ChIP-Enrich) or count based method (Poly-Enrich). This suggests that the regulation of certain processes is more often modified by multiple binding events (count-based), while others tend to require only one (binary). Our new hybrid method handles this by automatically choosing the optimal method, with correct FDR-adjustment.

**Author Summary:** Although every cell in our body contains the same DNA, our cells perform vastly different functions due to differences in how our genes are regulated. Certain regions of the genome are bound by DNA binding proteins (transcription factors), which regulate the expression of nearby genes. After an experiment to identify a large set of these regions, we can then model the association of these regions with various cellular pathways and biological processes. This analysis helps understand the overall biological effect that the binding events have on the cells. For example, if genes relating to apoptosis tend to have the transcription factor, Bcl-2, bind more often nearby, then Bcl-2 is likely to have a vital role in regulating apoptosis. The specifics of how to best perform this analysis is still being researched and depends on properties of the set of genomic regions. Here, we introduce a new, more flexible method that counts the number of occurrences per gene and models that in a sophisticated statistical test, and compare it to a previous method. We show that the optimal method depends on multiple factors, and the new method, Poly-Enrich, allows interesting findings in scenarios where the previous method failed.

## Introduction

Regulatory genomics experiments help us understand how cells use more than their genetic sequence to carry out a vast repertoire of cellular programs. Common regulatory genomics methods include chromatin immunoprecipitation followed by high-throughput sequencing (ChIP-Seq) and ATAC-seq, which identify transcription factor (TF) binding sites and open chromatin regions, respectively, across the genome. Other types of data, such as DNA Methylation assays, copy number alterations, repetitive element families and groups of SNPs, also lead to large sets of genomic regions that potentially play a specific role in regulatory genomics, with each type having notably different properties in terms of the number, size, and location of genomic regions.

Proteins that bind near a gene can regulate it in ways such as improving structural properties or physically blocking other proteins, often positively or negatively regulating the gene’s expression, respectively. Additionally, some proteins bind DNA several times in a clustered region [1], or in distant enhancer regions that interact with the same or distinct proteins bound in promoter regions [2]. Binding sites also differ in strength; a protein may bind in only a portion of cells in a sample at the time of immunoprecipitation, either due to weak binding or due to varying chromatin accessibility among the cell types in the sample. These binding sites along the genome are interpreted as peaks of varying strengths, depending on the signal-to-noise ratio or significance level of the peak. In general, interpreting each peak’s target gene(s) and effects remains an active area of research, which over time may improve results on downstream tests such as gene set enrichment.

Gene Set Enrichment (GSE) is an approach to test for over (or under) representation of genes in a set of genes with similar functionalities. Gene Ontology [3], Reactome [4], KEGG pathways [5], and MsigDB [6] are widely used gene set databases. Although originally developed for gene expression data, GSE testing is now often used to help interpret ChIP-seq peak sets and other sets of genomic regions. Existing methods for general GSE tests include Fisher’s exact test, random sets, logistic regression like LRPath [7], and GSEA-type tests [8]. GSE methods specifically for ChIP-seq data include Genomic Regions Enrichment of Annotations Tool (GREAT) [9], ChIP-Enrich [10], and Broad-Enrich [11]. With so many different tests, one may wonder which test is optimal to use, but there is no single recommendation across data types. Different tests are needed for different types of genomic regions as properties such as peak widths, number of peaks, and location relative to genes all make a difference. Thus GSE testing for genomic regions should not be a one-size-fits-all test; some methods work better than others in specific scenarios. For example, Cavalcante et al. showed that Broad-Enrich is more powerful than ChIP-Enrich for broad regions, but lacks power for narrow regions [11]. As another example, GREAT does not account for variability among genes, so it is best used in situations where the probability of a peak is constant across genomic space (e.g. per kb), as opposed to clustered near transcription start sites or displaying variability among gene loci.

Our previous method, ChIP-Enrich, uses a binary score to classify a gene as having at least one peak. We saw that ChIP-Enrich tends to underperform when nearly all genes have at least one associated genomic region; in this case, ChIP-Enrich will not yield meaningful results. We hypothesized that a count-based method that captures the frequency of binding would perform better in those situations. In this paper, we introduce such a method, Poly-Enrich to expand our available methods to be suitable for any set of narrow genomic regions including those that tend to saturate genes. Whereas ChIP-Enrich has the hypothesis that a single binding site is sufficient for regulation, Poly-Enrich allows for regulation that is incremental, i.e. more genomic regions correspond to stronger or more likely regulation. To identify under which situations one is more appropriate than the other, we performed a comparison of Poly-Enrich and ChIP-Enrich using a set of 90 transcription factor (TF) ChIP-seq datasets from the Encyclopedia of DNA Elements (ENCODE) [12]. We also introduce a hybrid test that combines information from both ChIP-Enrich and Poly-Enrich.

To illustrate the usage of Poly-Enrich, we apply it to sets of repetitive elements in the human Alu and LINE1 families, revealing for the first time a comprehensive view of the processes and functions enriched or depleted with these repetitive elements in the human genome. Finally, we describe several updates to our ChIP-Enrich website and *chipenrich* Bioconductor package, including additional methods for assigning genomic regions to target genes, new gene set databases, and more supported species.

## Results

### Motivation for development of Poly-Enrich

The motivation for our new methods comes from situations observed with real sets of genomic regions, often with ChIP-seq peak datasets, but also from other sources, such as families of repetitive elements or large sets of DNA polymorphisms such as those different between closely related species or sub-species. Although our original method, ChIP-Enrich, performs extremely well for most transcription factor (TF) ChIP-seq datasets (Figure 1a), because it uses a simple binary score for each gene, there are some scenarios where this simplification has a significant loss of information. For example, ChIP-Enrich models a gene with many peaks the same as a gene with only one peak, even though gene regulation may be affected by additional peaks (Figure 1b). Alternatively, if nearly every gene is assigned at least one peak, ChIP-Enrich would be unable to distinguish among them and thus unable to detect any gene set enrichment (Figure 1c). We therefore developed Poly-Enrich as a count-based method that addresses both of these scenarios.

**Figure 1:**
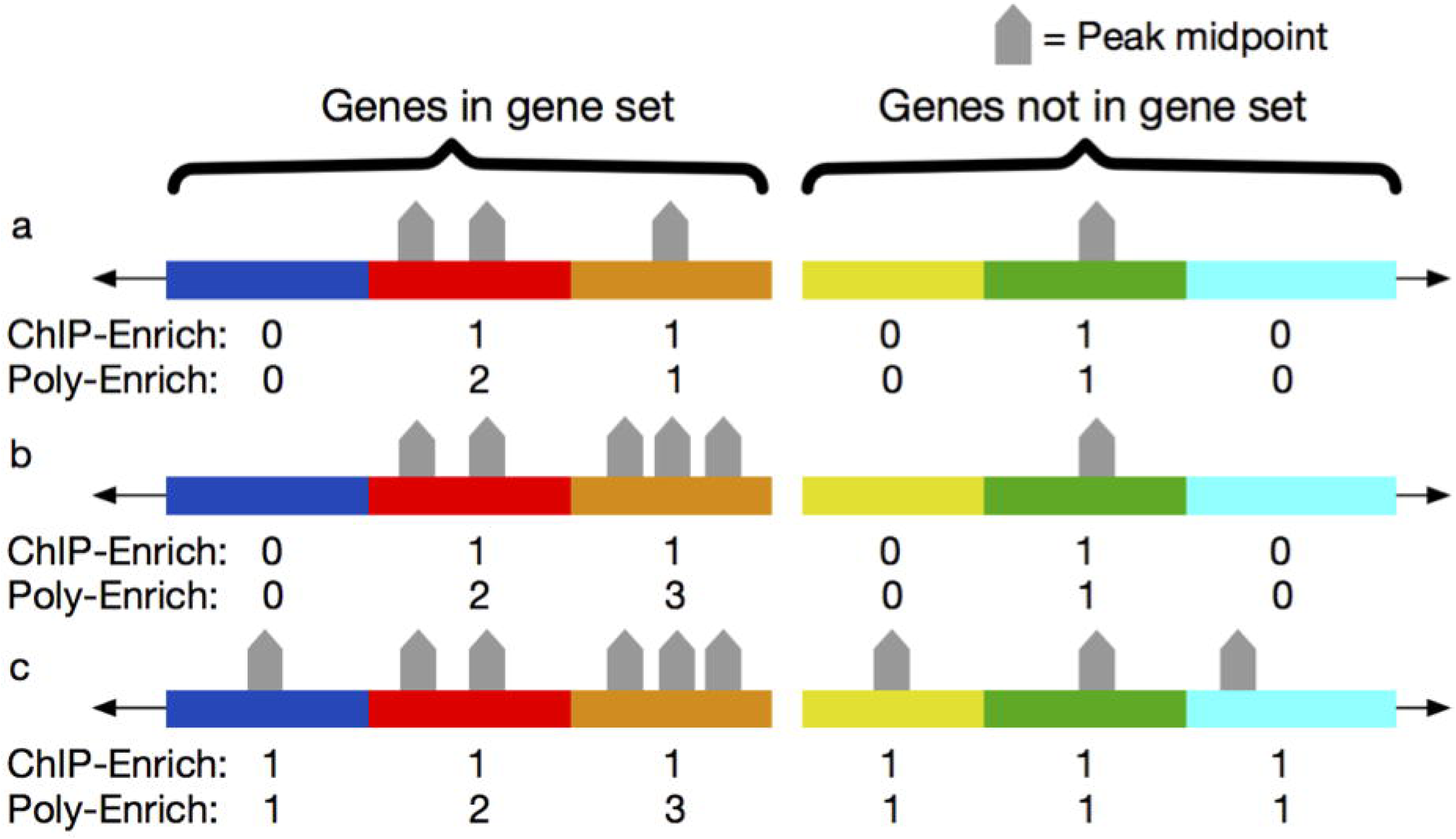
Three scenarios of ChIP-seq peak distributions illustrating how ChIP-Enrich and Poly-Enrich perform. Each color represents a different gene locus; the left three are in a gene set and the right three are not. (a.) Peaks are relatively evenly distributed, with a small number across a subset of genes. Given this situation, ChIP-Enrich evaluates 2/3 vs 1/3 while Poly-Enrich evaluates {0,2,1} vs {0,1,0}; both methods perform well. (b.) Some genes contain significantly more peaks than others, such that information is to be gained from the number per gene. ChIP-Enrich evaluates 2/3 vs 1/3, Poly-Enrich evaluates {0,2,3} vs {0,1,0}; ChIP-Enrich performs adequately, but Poly-Enrich is optimal. (c.) Nearly all genes have at least one peak, with some having significantly more than others. ChIP-Enrich evaluates 3/3 vs 3/3, Poly-Enrich evaluates {1,2,3} vs {1,1,1}; ChIP-Enrich would not detect any enrichment, while Poly-Enrich can still detect gene sets enriched with more peaks.

Although GREAT is also a count-based gene set enrichment method, Poly-Enrich differs significantly from it in two respects. Firstly, whereas GREAT counts the number of peaks in an entire gene set, Poly-Enrich counts them per gene. By separating counts per gene, we are able to adjust for each gene’s locus length and the variability in peak count across genes, which we previously showed was an important adjustment to control for Type I error [10]. Secondly, the binomial model used by GREAT assumes that the background probability of a peak is constant across the genome. Poly-Enrich uses a more flexible, empirical approach to this that provides for a range of different assumptions about peak distribution.

### Assigning peaks to genes

Genomic regions can be assigned to genes in different ways, so that regulation from different types of regions (e.g., promoters, introns, or regions distal to TSSs) can be studied. For example, ChIP-Enrich, GREAT, and Poly-Enrich all use a peak’s midpoint to define the location of the peak. We define a gene’s locus definition as the region on the genome such that peaks in that region are assigned to the gene. These loci are defined using properties of the gene, such as within 5kb of a gene’s transcription start site (TSS), or simply by assigning each region to the nearest TSS (Figure 2). In the new version of our website and Bioconductor package, we offer several additional choices, including exons, introns, and distal regions only (>10kb upstream from a TSS). Users can also upload their own custom locus definition, such as open chromatin regions for a specific cell type, or known enhancers and their target genes.

**Figure 2:**
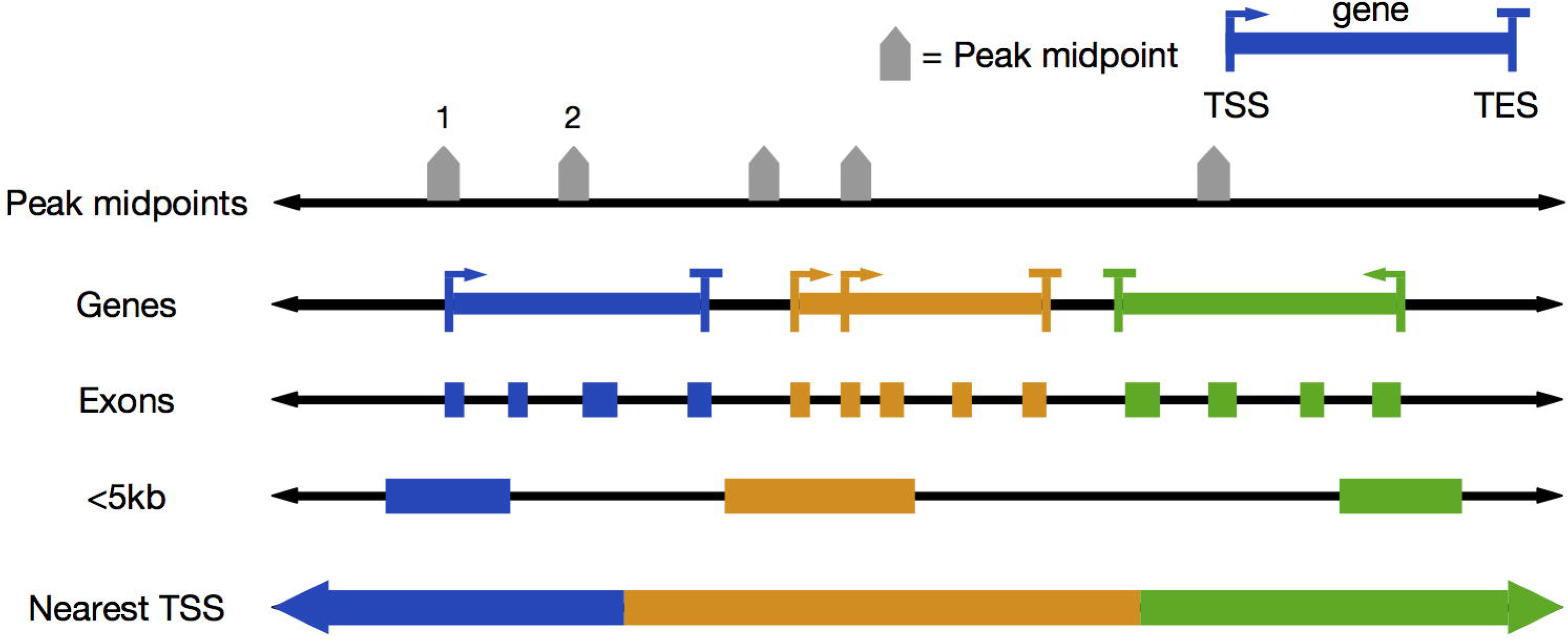
Overview of peak-to-gene assignments given gene locus definitions. Examples shown are: “<5kb”, peaks within 5 kilobases of a gene’s TSS are assigned to the gene; “Nearest TSS”, peaks are assigned to the gene with the closest TSS. A gene’s locus length is defined by the number of base pairs assigned to the gene. In this toy example, peak 1 would be assigned to the blue gene in the <5kb locus definition and for Nearest TSS, while peak 2 would not be assigned to any gene in the <5kb locus definition.

### Poly-Enrich model

We model the number of peaks per gene using a negative binomial generalized linear regression as a function of gene set membership, and with a cubic smoothing spline to empirically model the relationship with gene locus length:

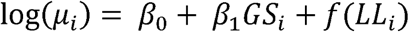

where for gene *i, μ*_*i*_ is the expected mean number of genomic regions assigned to the gene, *GS*_*i*_=1 if the gene is a member of the gene set of interest and 0 otherwise, and *f*(*LL*_*i*_) is a negative binomial cubic smoothing spline to adjust for the gene’s locus length. We then look at the sign and significance of *β*_1_ to test for enrichment, where a positive *β*_1_ indicates enrichment, and a negative value indicates depletion (fewer regions than expected at random).

### Testing Type 1 error and power

We tested the type I error rate of the count-based method under the null hypothesis of no enrichment signal. By permuting the genes in the peak-to-gene assignment pairs and breaking the peak-gene relationships, we simulated three scenarios of no enrichment: *i)* the *“complete”* randomization was done by shuffling the gene IDs in the whole dataset; *ii)* the *“bylength”* randomization first groups the genes together into bins of similar locus length, and then randomizes genes within those bins to preserve the locus length relationship; *iii)* the *“bylocation”* randomization groups genes together by their physical location on the chromosomes, and then randomizes genes within those bins. (See *Methods* for more detail.) We ran the randomizations on our 90 selected ChIP-Seq datasets from ENCODE (see *Methods*), 10 times each, and the proportion of p-values < 0.05 and < 0.001 for each dataset were plotted (Supplementary Figure 1). We see that the test is properly controlled at an acceptable level for Type 1 error in all cases.

To characterize the statistical power of Poly-Enrich under different situations, we permutated data while simulating enrichment of a gene set, and compared the results with those from ChIP-Enrich. We used three datasets with a small, medium, and large number of peaks, and two GO terms with a small and large number of genes. Three types of enrichment were simulated: one that adds peaks mainly according to the regulatory assumptions of ChIP-Enrich (CEBias), one that adds peaks mainly according to the assumptions of Poly-Enrich (PEBias), and one that is balanced. For each type of enrichment, we simulated four levels of enrichment: 0.05, 0.1, 0.2, and 0.3, which indicates the proportion of additional peaks added to the gene set. (See *Methods* for more detail.) Finally, we chose two different levels of significance: *α* = 0.05 and 0.001, as our cutoffs.

As expected, a larger gene set and higher simulated enrichment results in higher power. Overall, we see that Poly-Enrich has more power in simulations that enrich a gene set by increasing the number of peaks per gene, and ChIP-Enrich has more power than Poly-Enrich in simulations that enrich a gene set by adding peaks to genes without them. Finally, the *Balanced* simulation results in the two methods having similar power in most cases (Supplementary Figure 2). Simulations on larger datasets artificially reduced power because randomized larger datasets include more noise. However, in real experiments, we expect larger datasets (more peaks) to be more powerful, since the majority of the peaks in the dataset are not expected to be noise.

### Poly-Enrich with weighted genomic regions

The height and confidence of peaks in a ChIP-seq experiment can vary dramatically, and we reasoned that incorporating this additional information would improve the ability to pinpoint the truly enriched pathways. Although the most apparent motivation for weighting genomic regions is to account for ChIP-seq peak strength, there are other situations where each peak or genomic region may be assigned a unique score (e.g. confidence or strength). Therefore, we added the option to weight regions by signal value (see *Methods* for details), and examined the extent to which adjusting for peak strength improves enrichment results using 90 ENCODE ChIP-seq datasets by comparing the −log_10_ p-values per gene set. To also ensure that no enrichment result flipped from enriched to depleted or vice versa, we used a signed −log_10_ p-value, where values for depleted gene sets were negative, and values for enriched gene sets were positive.

We noticed for 25% of the experiments, most enriched gene sets were more significant with weighting, thus as we hypothesized, binding events near genes in enriched GO terms were stronger than those near other genes (Figure 3A, 3B). In another 20% of the experiments, the enrichment p-values were split between the two methods (Figure 3C). Interestingly, the distribution of log signal values for these experiments showed a bimodal pattern (Figure 3D). This suggests that some gene sets tend to have genes with significantly stronger binding peaks than others, and that both sets may be biologically interesting. For the remaining 55% of experiments tested, weighting made little difference on the results.

**Figure 3:**
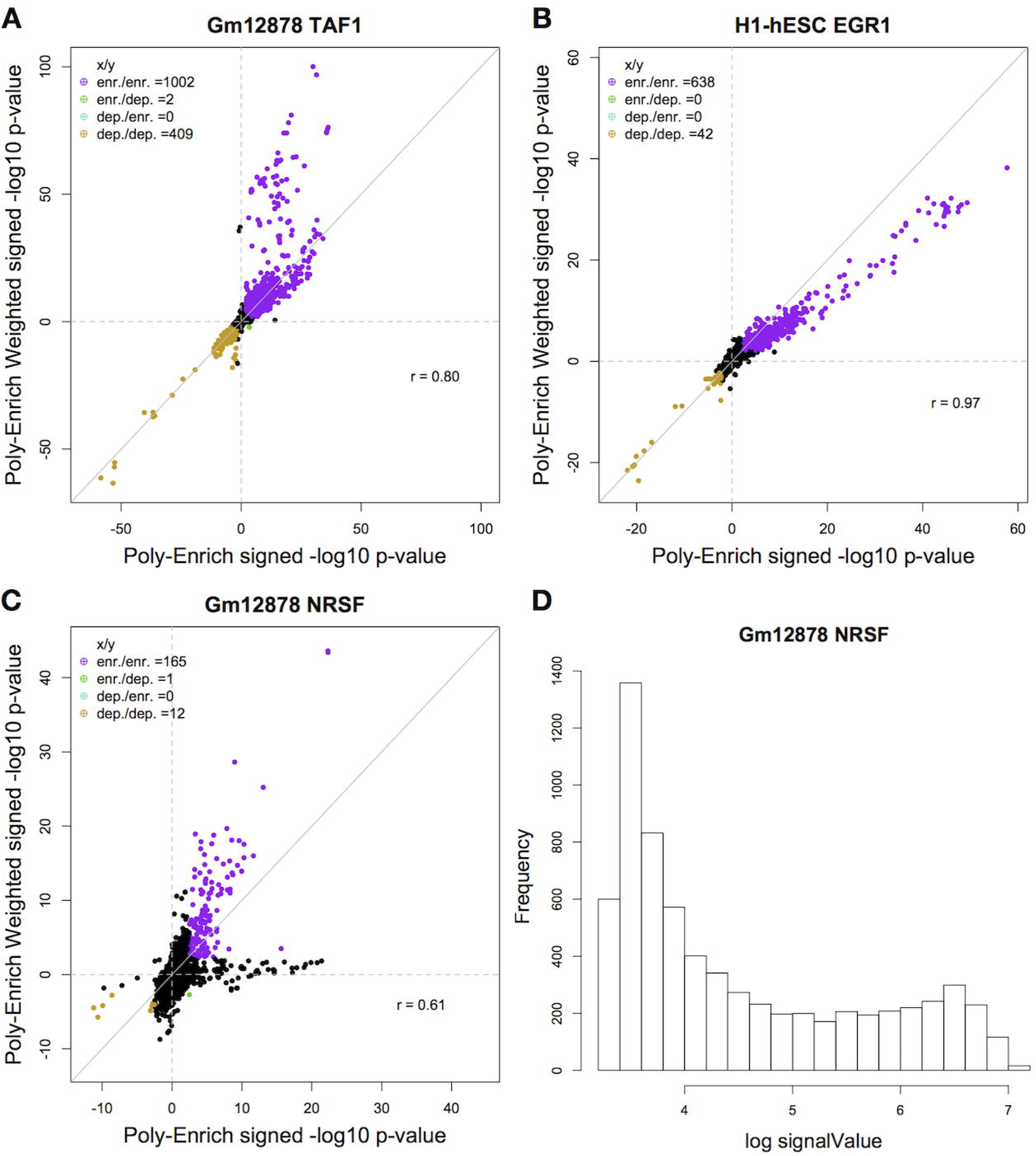
Comparison of GO term enrichment results between standard Poly-Enrich and its weighted version using signal values as weights. Each point is a GO term’s −log_10_ p-value of the two methods, signed positive for enriched, negative for depleted. (A) Using weighting results in more significant enrichment in many GO terms in the Gm 12878 TAF1 ChIP-Seq experiment. (B) Using weighting results in less significant enrichment in many GO terms in the H1-hESC EGR1 ChIP-Seq experiment. (C) Using weighting on the Gm 12878 NRSF experiment results in several more significant GO terms as well as several less significant ones. (D) The histogram of log signal values from the NRSF experiment. There is a bimodal pattern in the weights, suggesting that there are some GO terms with genes that tend to have stronger or weaker binding.

### Comparison of the count-based (Poly-Enrich) versus binary (ChIP-Enrich) model of enrichment

We next compared signed log10 p-values from Poly-Enrich versus ChIP-Enrich on the same set of 90 ENCODE ChIP-seq datasets. Our initial hypothesis was that some experiments would be clearly modeled better by one method or the other (i.e. dependent on the transcription factor). However, our results strongly suggest that the optimal method is more dependent on the gene set than the TF. This can be visualized by a split in the significance levels of GO terms between the binary and count-based methods (Figure 4A), and suggests that a single transcription factor may regulate genes differently depending on the function of the gene. Thus, we sought to understand this further.

**Figure 4:**
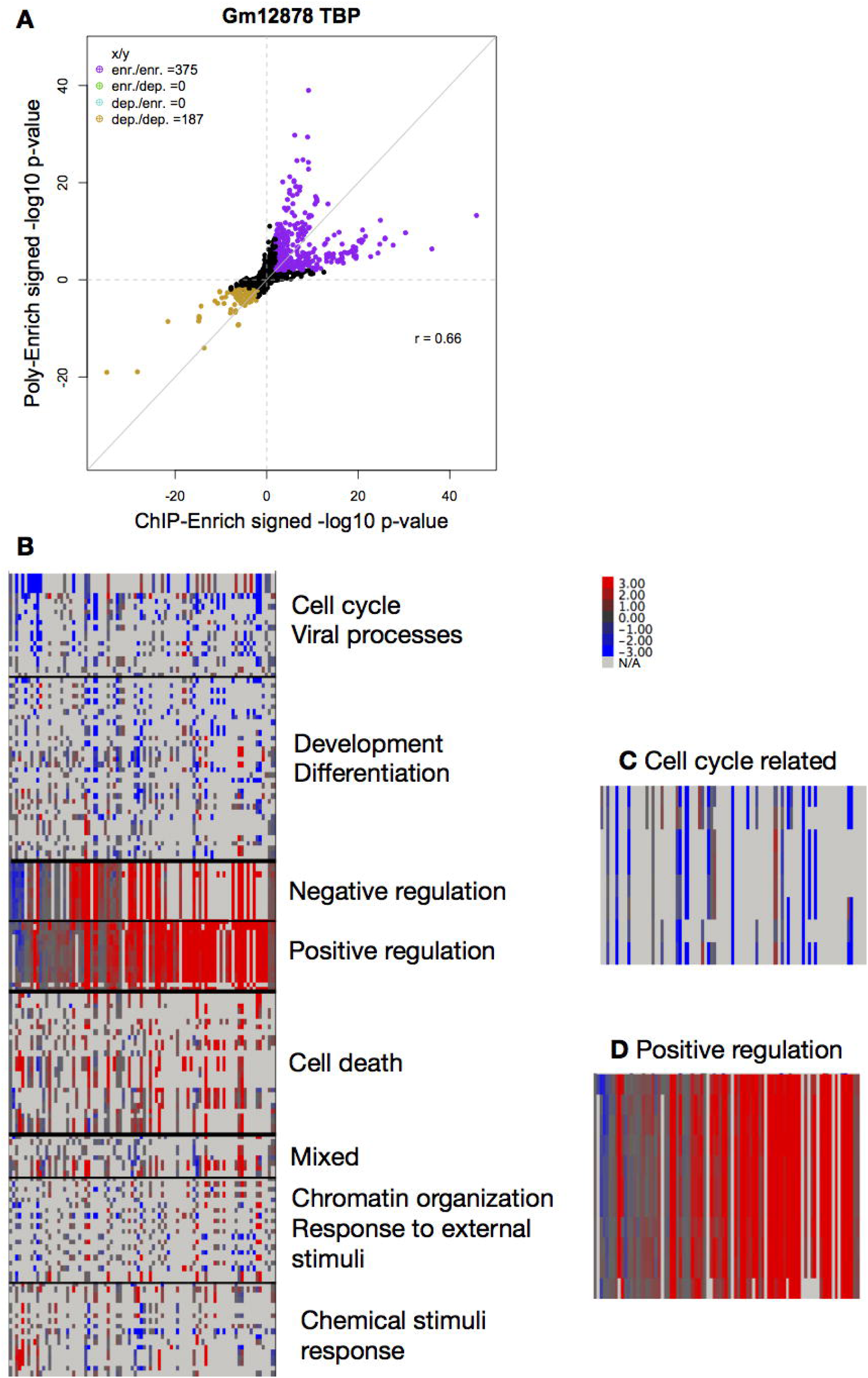
(A) Comparison of GO term significance levels between ChIP-Enrich and Poly-Enrich. Each point is the –log_10_ p-value of a GO term from the two methods, signed positive for enriched or negative for depleted. Several gene sets are much more significant using ChIP-Enrich and several are much more significant using Poly-Enrich, however 32% of the datasets showed a split pattern like shown. (B) Heatmap of −log_10_ p-value differences between Poly-Enrich and ChIP-Enrich for GO terms and ChIP-seq experiments, where each row is a GO term and each column is a ChIP-seq experiment. Shown are GO terms where more than 15% of the experiments had a −log_10_ p-value difference of 2 or larger. Red indicates Poly-Enrich was more significant, and blue indicates ChIP-Enrich was more significant. Light grey indicates the transcription factor used in the experiment was not assigned to the GO term and is omitted in the clustering. Representative GO terms are shown for each cluster. (C) GO terms related to cell cycle are mostly blue, indicating that a binary score provides a more appropriate model. (D) GO terms containing “positive regulation” are mostly red, indicating that a count score provides a more appropriate model.

The binary model used by ChIP-Enrich assumes that a single binding event (i.e. a single genomic region) is sufficient for regulation, while the Poly-Enrich count-based model assumes that strength of regulation is incremental with the number of binding sites. Based on the results above, we asked what kinds of genes were more consistent with either of those assumptions. To answer this, we first created a set of true positives comprised of GO term-TF pairs by using the GO term biological process (BP) assignments for the gene encoding the transcription factor (e.g. the gene encoding for JunD is assigned to the GO term, “cell death”). This gold standard makes the reasonable assumption that TFs tend to regulate genes in the same biological processes in which they are active. Observing the enrichment results using the 5kb locus definition for these true positive GO term-TF pairs, we used clustering to identify patterns of TFs and GO terms that are optimal with one of the methods. We found that the method that worked better was most often determined by the GO term (Figure 4B). For example, GO terms related to “cell cycle” clustered together and displayed greater power with ChIP-Enrich. Conversely, GO terms involving “positive regulation” tended to do better with Poly-Enrich except for those involving cell cycle (Figure 4C,4D). Parallel results using the Nearest TSS locus definition were similar (Supplementary Figure 3).

Since the gene set, rather than the experiment, was a stronger determinant of the more appropriate method, in many cases we are unable to recommend either Poly-Enrich or ChIP-Enrich for an entire experiment; one exception is that Poly-Enrich is recommended for experiments with a very large number (>100k) of peaks. We therefore developed a hybrid test that uses information from both ChIP-Enrich and Poly-Enrich.

### Hybrid test

To obtain the best results across all types of GO terms and datasets, we developed a hybrid test that incorporates both the binary and count-based models. After performing the two models, the hybrid p-value of the two tests is defined as: *P*_*hybrid*_ = 2 × min (*P*_*CE*_,*P*_*PE*_) [13], where *p*_*CE*_ and *p*_*PE*_ are the p-values given by ChIP-Enrich and Poly-Enrich, respectively. This is essentially a Bonferroni-adjusted p-value for two tests. This hybrid has been shown to be beneficial if the two tests are sufficiently different, but loses power and is conservative if the tests are identical or nearly identical [13]. While the hybrid test is not as powerful as the better method between ChIP-Enrich and Poly-Enrich, it is dramatically more powerful than using the worse method, making it the optimal method to use across all GO terms (Figure 5). While this hybrid test currently only accommodates ChIP and Poly-Enrich, we can extend this to accommodate several additional gene set enrichment tests.

**Figure 5:**
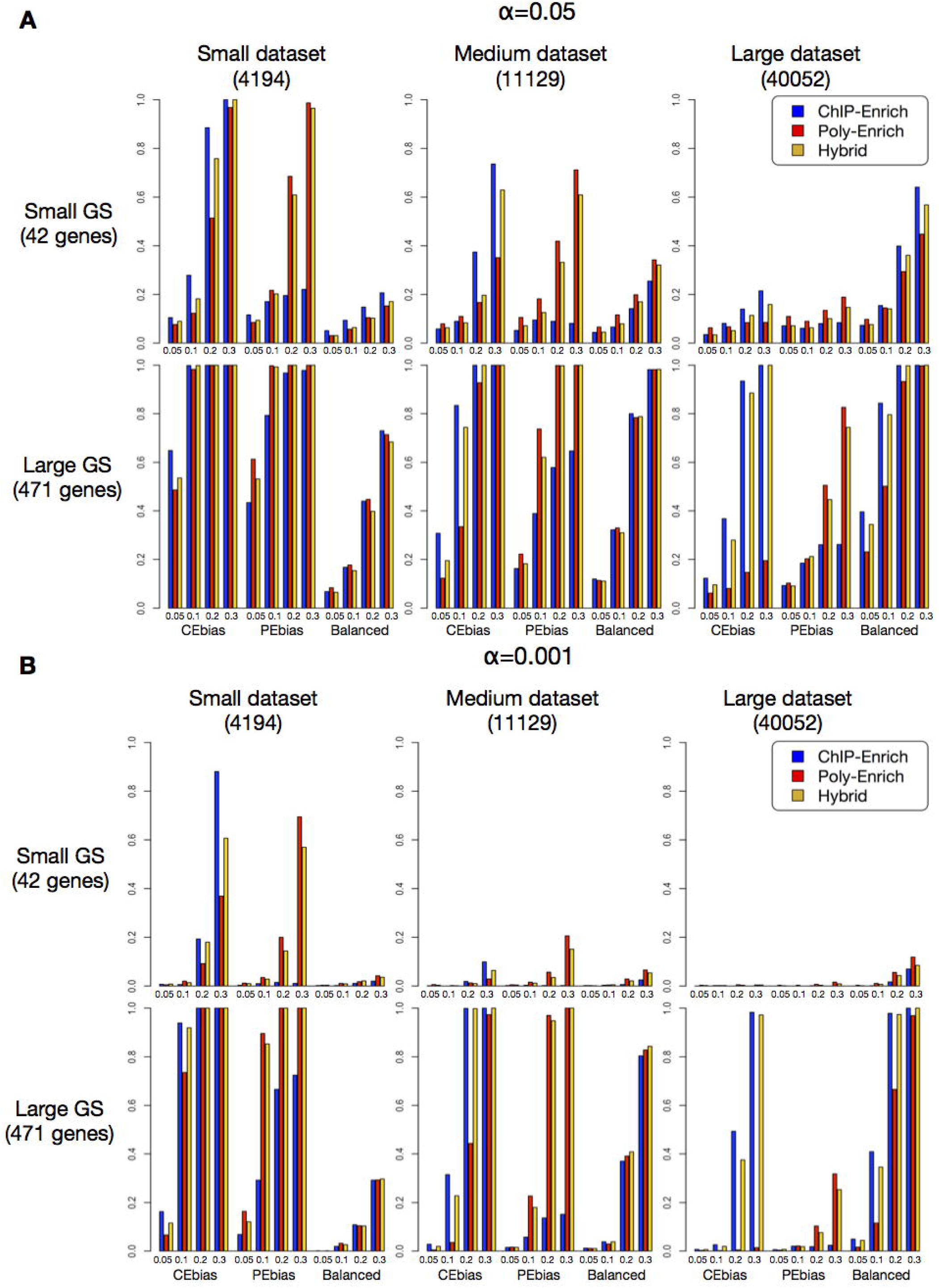
Statistical power comparisons for Poly-Enrich (red), ChIP-Enrich (blue), and the hybrid test (gold) for datasets with three different sizes (i.e. number of peaks: small, medium, and large) and two gene set sizes (small and large GS), under two significance levels: *α* = 0.05 (A) and 0.001 (B), and three different methods of simulated enrichment (CEbias: add peaks according to the regulatory assumptions of ChIP-Enrich, PEbias: add peaks mainly according to the assumptions of Poly-Enrich, Balanced: add peaks proportional to each gene’s locus length). The values on the X-axis indicate the percent of extra peaks added to simulate enrichment; a higher value simulates stronger enrichment. The hybrid test is shown to have much more power than the wrong method, but only a bit less power than the correct method.

### Identifying biological processes enriched with or depleted in repetitive element families using Poly-Enrich

To further illustrate the utility of Poly-Enrich, we used it to test sets of repetitive element regions. We asked whether we could identify gene sets that tended to be either enriched or depleted for certain types of repetitive elements. Significant enrichment of the repetitive elements in the promoter regions of genes, for example, can sequester the transcription factors that will inhibit activities at another transcription factor binding site or other regulatory motif [14]. Some of these mobile elements remain active with new insertions having neutral, detrimental, or beneficial effects. Although repetitive element families have been well studied for over 30 years, little is yet known about the biological processes that they have adapted to help regulate or that they can too easily disrupt and thus are negatively selected against [15]. Using the database of human repetitive elements from the UCSC Table Browser (RepeatMasker 3.0) [16], we performed GSE testing on repetitive element families. Certain families of repetitive elements have over a million occurrences across the human genome, and thus virtually all genes have at least one nearby instance, making this an example where ChIP-Enrich performs poorly. Thus, in this situation, modeling the number of insertions per gene is critical to identify differences.

We examined two of the most abundant types of repetitive elements: the *Alu* and LINE1 (L1) elements, which make up an estimated 11% and 17% of the human genome, respectively [17, 18]. We also chose four gene locus definitions: Nearest TSS, <5kb (promoter regions), >5kb (distal regions), and Intron. We tested GO Biological Processes, and used hierarchical clustering of the resulting GSE significance levels to identify related groups of biological processes enriched with or depleted of the repetitive elements (Figure 6). We found strong enrichment of Alu’s in GO terms describing metabolic processes, most significantly “ATP metabolic process” and “rRNA metabolic process”, especially in promoter regions, which is consistent with an analysis of *Alu* distribution in chromosomes 21 and 22 that showed *Alu* elements on these chromosomes were enriched in or near metabolism and signaling genes [19]. Conversely, Alu elements were sharply depleted in the promoter regions of many development and morphogenesis gene sets, with the strongest depletions in “cell fate commitment” and “connective tissue development”. Interestingly, depletions were also seen in the introns of genes in these gene sets, but not in regions >5kb upstream, suggesting the negative selection (depletion) is limited to the regions that are more commonly regulatory.

**Figure 6:**
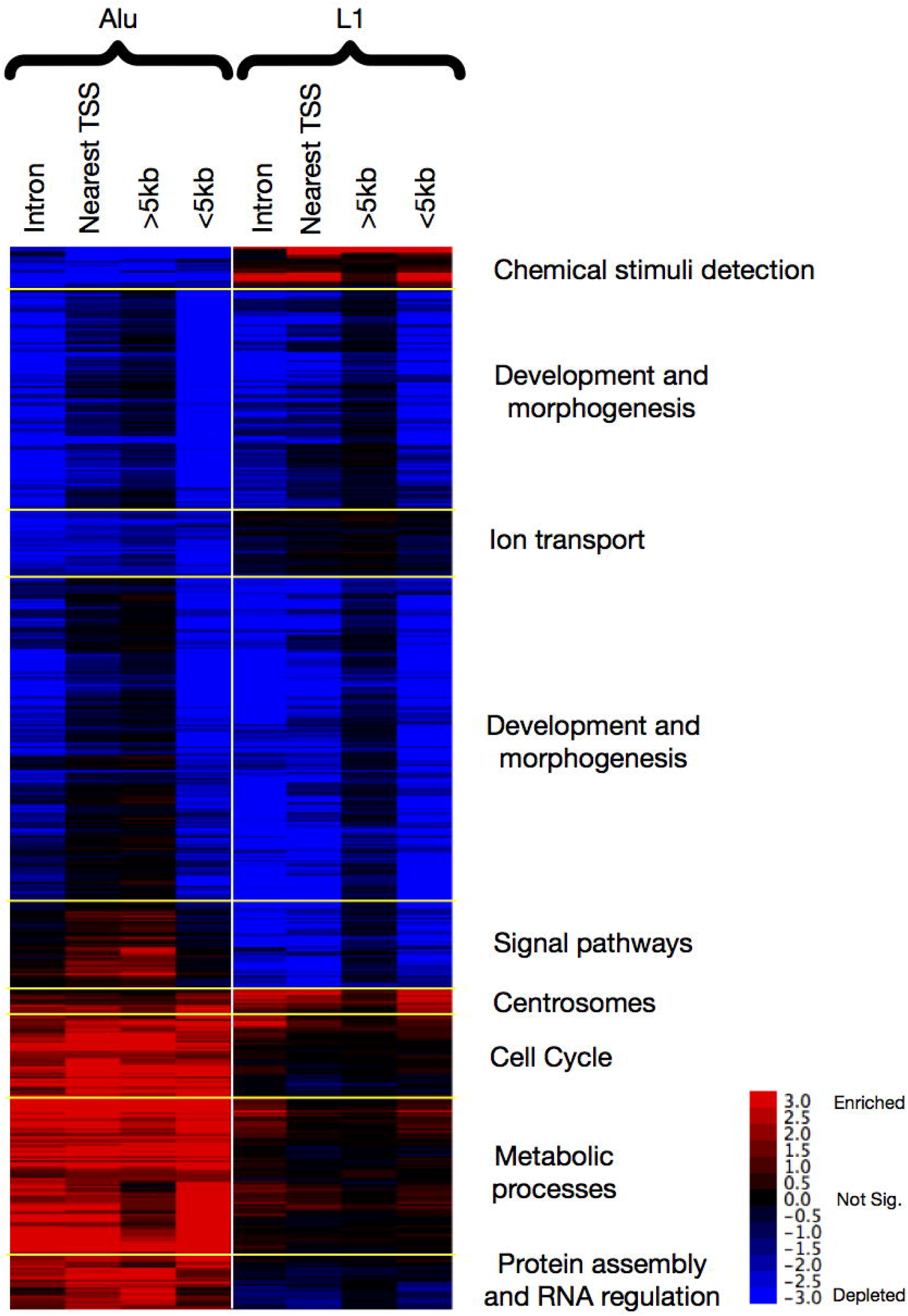
Enrichment results using Poly-Enrich for Alu (first four columns) and L1 (last four columns) repetitive element families using four different peak-to-gene assignments. Shown are signed −log_10_ FDR, where positive values (red) indicate enrichment and negative values (blue) indicate depletion. Only GO terms that were significant for at least 3 columns at the FDR = 0.05 level are displayed. We identified nine clusters of GO terms with similar enrichment patterns. Representative GO terms are used to label each cluster.

Novel insertions of L1 elements into or near key genes are known to be associated with neurological diseases [20]. Consistent with this, we found that all neuro-related GO terms in Figure 6 were depleted for L1 (but not for Alu) (Supplementary Figure 4), which suggests that L1’s evolutionarily have been selected against occurring in the regulatory regions of neurological genes; when they are inserted into the introns or promoters of these genes, the inserted elements may cause disease.

In general, we observed that the significance of the distal upstream regions (>5kb locus definition) was lower than the other three locus definitions (with the exception of some enrichments for Alu elements) (Supplementary Table 1), implying that most repetitive element negative (or positive) selection has occurred in the promoter regions or introns of genes. Alternatively, the gene distal enriched and depleted regions may be limited to a specific set of enhancer regions, the signal from which could have been diluted in our analysis. Interesting additional findings are that L1 elements are enriched in chemical stimulus detection such as “detection of chemical stimulus” and “sensory perception of chemical stimulus”, while Alu elements are depleted in the genes in these processes. We also find that both Alu and L1 are enriched in centrosome-related GO terms, which were only made possible with recent advancements in genome mapping near the centromeres [21], and is consistent with previous findings [22]. Finally, both Alu and L1 elements are significantly depleted in genes in GO terms relating to development and morphogenesis.

### Availability, usage, and updates

Poly-Enrich is available in the *chipenrich* Bioconductor package and as a web interface at http://chip-enrich.med.umich.edu. Several additional gene set databases and gene locus definitions have been added for the user to choose (see http://chip-enrich.med.umich.edu/data/ChipenrichMethods.pdf).

To perform gene set enrichment analysis, the user first needs a file of genomic regions, which may be a narrowPeak, BED, or text file with chromosome, start, and end positions for each region. The user then selects a species, one or more gene set databases, a gene locus definition, and the test method (ChIP-Enrich, Poly-Enrich, Hybrid, or Fisher’s exact test); the gene set and locus definition can be built-in or user-defined. The user can then also choose to add weight based on a peak-specific score, and a number of other options, such as adjustment for read mappability.

The enrichment function outputs five files:

- opts: The options that the user input into the function.
- peaks: A peak-level summary showing the peak-to-gene assignments for each peak.
- peaks-per-gene: A gene-level summary showing gene locus lengths and the number of peaks assigned to them.
- results: The results of the GSE tests. Lists the tested gene sets along with their descriptions, the test effect, odds ratio, enrichment status, p-value, and FDR. Also included is the list of gene IDs with contributing signal for each enrichment test.
- qcplot: A plot of the gene locus lengths with a fitted smoothing spline.

## Discussion

Gene set enrichment testing methods for genomic regions should take into account the differing properties of the input datasets, including the widths and number of genomic regions, and where they tend to occur relative to genes. However, no single method is appropriate for all types, and therefore no single GSE method should be recommended for all sets of genomic regions. Although our previously developed ChIP-Enrich method for gene set enrichment with genomic regions performs well for most transcription factor ChIP-seq datasets [10], above we described some common situations where it does not. Such cases include when nearly all genes are assigned at least one genomic region, and when the strength or likelihood of regulation increases incrementally with the number of genomic regions. As an example, the transcription factor NF-kappaB is known to regulate the gene NFKBIA by binding to a few or even many motif positions in the promoter [23], with gene expression correlated with the number of bound factors. Thus, motivated by specific examples of regulatory mechanisms, we developed Poly-Enrich, a method that models the number of regions per gene, empirically adjusts for each gene’s locus length, and takes into account variability among genes in each gene set. Poly-Enrich is also flexible, in that it easily allows for weighting of each genomic region by any score of interest. We used the example of weighting by peak strength, but other examples include weighting by SNP significance in a GWAS analysis, by the inverse distance to a gene, or by the probability that the region is in an open chromatin region in a particular cell type.

We showed that our count-based method, Poly-Enrich, works well when almost all genes are assigned a peak, whereas ChIP-Enrich does not. In comparing when each is most appropriate, we discovered that it is mostly dependent on the gene set, rather than the transcription factor. Because in many cases we can not recommend a single best method to test all gene sets for an experiment, we developed and implemented a hybrid test that uses information from both methods and performs better than either test across GO terms for most datasets.

When applying Poly-Enrich to repetitive element families, we both reconfirmed known associations and also identified novel findings. Poly-Enrich confirmed that Alu elements tend to regulate genes for metabolism and signaling by finding enrichment for related GO terms. Additionally, we know that L1 insertions into or near certain neurological-related genes are associated with neurological diseases [24]. We find that L1 is depleted in neuro-related GO terms, implying there normally are fewer L1 elements in the regulatory regions of these genes, which is consistent with neurological diseases being associated with L1 element insertions near these genes. We also find that there is little enrichment or depletion in the distal regulatory regions of genes, suggesting that repetitive elements may not have as large of an affect there due to mitigated regulatory activity at larger distances from transcription start sites. Poly-Enrich also detected some novel associations between repetitive element families and biological pathways. Both Alu and L1 elements were significantly depleted in genes in GO terms relating to development and morphogenesis, such as “connective tissue development” and “skeletal system morphogenesis”, suggesting that it is especially critical to have developmental regulatory regions free from potentially disruptive repetitive elements during early growth.

One shortcoming of our current methods (as well as current alternatives) is that they rely on associating each genomic region with the nearest gene. However, it is estimated that 79-95% of DNAse I hypersensitive sites, markers for enhancer regions, actually regulate a different, distal target gene [24, 25]. We are currently developing a set of enhancer locus definitions that identify and assign enhancer regions to their appropriate target genes, as was recently introduced by Chicco D, et al [25], so peaks in enhancer regions will be correctly assigned and false positive peaks in nonfunctional intergenic regions will be filtered out. We believe this will improve all future gene enrichment analyses.

## Methods

### Datasets

All ChIP-Seq data were obtained from Encyclopedia of DNA Elements (ENCODE) at University of Califonia, Santa Cruz [12]. We chose a total of 90 experiments over the three Tier 1 cell lines (Gm12878, H1-hESC, and K562), and all 35 transcription factors that had available ChIP-seq data for at least two of the three Tier 1 cell lines. (Supplementary Table 2)

The gene sets used were from Gene Ontology: Biological Processes (GOBP) ver. 3.4.2 [3]. We filtered out gene sets with less than 15 genes or more than 2000 genes as gene sets with too few genes generally have insufficient power and may not satisfy the assumptions of the statistical model, and gene sets with too many genes are too vague to be biologically informative. In total, there were 5015 gene sets.

### Assigning regions to genes

The UCSC knownGene database for hg19 was used to define the transcription start sites across the genome [26]. Each locus definition (e.g. nearest TSS, <5kb) was generated as a table containing the columns: chromosome, Start, End, gene ID. All genomic regions whose midpoint was between a gene’s start and end values would be assigned to that gene. It is possible for a locus definition to have several disjoint regions for a certain gene.

### Poly-Enrich model: a generalized linear model with a negative binomial family

We model the number of genomic regions assigned to each gene with a generalized linear model (GLM) with a negative binomial (NB) family. The model is:

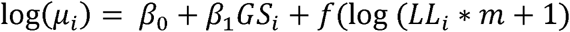

where for each gene *i, GS* is an indicator for whether the gene is in the gene set of interest or not (=1 if in the gene set; 0 otherwise), *μ.* is the mean of the negative binomial distribution for the number of genomic regions assigned to each gene, and the overdispersion parameter *θ* is estimated so that *Var*(*Y*|*GS*) = *μ* + *θμ*^2^, where Y is the number of genomic regions for the gene. The function *f* is a negative binomial cubic smoothing spline that adjusts for the gene’s locus length and optionally adjusts for *m*, the mappability of the gene’s locus. Details about mappability can be found in the ChIP-Enrich manuscript [10]. We use the *gam* function in the *mgcv* R package to fit the model, which uses a penalized likelihood maximization, and the smoothing spline penalty is a squared second derivative penalty [27]. Use of a cubic smoothing spline to adjust for the genes’ locus lengths was first introduced in ChIP-Enrich, as a powerful, flexible way to model this relationship.

A Wald test on the coefficient for the gene set is used to test for enrichment (or depletion) of each gene set: the test statistic is defined as 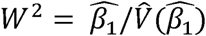, which follows a 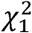 distribution under the null hypothesis that there is no association between gene set membership and number of genomic regions.

### Poly-Enrich with weighting based on genomic region scores

In certain cases, each genomic region in a dataset may be associated with a numeric score. For example, ChIP-seq peak finding results often include a value denoting the strength of a peak, (e.g. signalValue in ENCODE dataset results or −10*log_10_(p-value) in MACS2 results). Poly-Enrich weights based on these scores by giving each genomic region a weight proportional to its signal value (or other score) and normalizing such that the mean of all weights is equal to 1. For every genomic region assigned to a gene, we sum all the weights and substitute the weighted sum in place of the original count. The same model is used, except assuming a quasi-negative binomial family to accommodate for non-whole number data. The calculations can be carried out identically to the standard negative binomial family.

### Comparing p-values between methods

To compare p-values between methods, we use a scatterplot, plotting a signed −log_10_ p-value per gene set. If a gene set is enriched, the sign is positive, and if the gene is depleted, the sign is negative. This allows us to detect if there are any cases where two methods may contradict each other’s conclusions.

### Spline approximation for Poly-Enrich

With a library of over 20,000 genes and most gene sets being less than 1000 genes, the cubic smoothing spline estimate changes very little between gene sets. Thus, we have confirmed we can reasonably assume that the spline is approximately equal for any gene set of interest, including the spline with no gene set (Supplementary Figure 5A).

We first run the same model except without the gene set (*GS*) term:

Log(*μ*_*i*_) = *β*_0_ + *f*(*LL*_*i*_). We then extract the fitted spline using the *predict* function with *type=”terms”* from the *mgcv* R package to obtain a spline-adjusted locus length for each gene. This new value is then input in the model for every gene set, which allows us to fit a spline only once instead of once for each gene set. This saves a significant amount of time when testing a large number of gene sets (approximately 75% time saved when testing 4000 gene sets). Compared to the original model, we find that the p-values from the spline approximation model are nearly identical (Supplementary Figure 5B, 5C).

### Use of Score test for quick approximations

One of the alternatives for the Wald test is the Score test [28]. To obtain quick, approximate enrichment results for many sets of genomic regions (e.g. a large group of ChIP-seq peak datasets) using either the Poly-Enrich or ChIP-Enrich approach, we now provide the Score test approximation for convenience. We can calculate the score test statistic for ChIP-Enrich as:

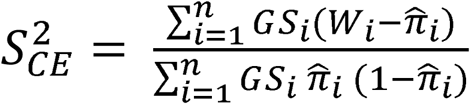

where for each gene *i, GS*_*i*_=1 if the gene is in the gene set of interest and 0 otherwise, *W*_*i*_=1 if the gene has at least one genomic region and 0 otherwise, and 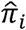 is the predicted probability of the gene having at least one genomic region obtained using the *fitted* function with *type=”terms”* in the *mgcv* package, assuming the null hypothesis of no association (i.e. *β*_1_=0 in the model).

The score test statistic for Poly-Enrich is:

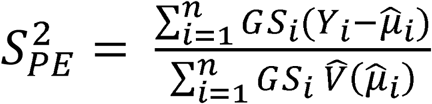

where for each gene *i, GS*_*i*_=1 if the gene is in the gene set of interest and 0 otherwise, *Y*_*i*_ is the number of genomic regions assigned to the gene, 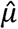 is the predicted number of genomic regions assigned to the gene obtained using the *fitted* function with *type=”terms”* in the *mgcv* package, assuming the null hypothesis of no association, and its variance is estimated empirically, assuming *GS*_*i*_ = 0 in the variance calculation.

The advantage of using the score test is that all the required parameters are already estimated during the spline approximation, so all subsequent calculations will no longer require fitting of a GLM. This reduces the runtime of the enrichments significantly further (approximately 95% time saved when testing 4000 gene sets). However, there are some scenarios where the Score test differs from the Wald test by a substantial amount, mostly notably for depleted gene sets, so the default option is set to the Wald test (Supplementary Fig 6). If results are desired quickly, the Score test is offered (method=”chipapprox” or “polyapprox”) and can serve as a convenient approximation.

### Testing Type I error

The null hypothesis of Poly-Enrich is that there is no true biological enrichment. To test the Type-I error, we randomly permute the genes to simulate a scenario where there is no association between genes and the number of peaks. However, to ensure that the results are not biased by gene locus length or gene location, we performed two additional permutations: one permutes genes within bins of similar locus length, while the other permutes within bins of chromosomal locations. In both cases, the genes are sorted by the variable of interest (locus length or location), and then assigned to consecutive bins of 100 genes each.

For each of the 90 TF peak datasets chosen, after assigning the peaks to genes, we permute the gene IDs using the randomization of interest, and then perform enrichment tests against GO biological processes. We ran a total of 10 trials and took the median p-value per gene set as the randomization p-value. Then, the proportion of p-values less than a defined confidence level was determined per experiment to calculate the overall Type I error. We then plotted all 90 overall Type I errors for each experiment in a box plot to convey overall Type I error.

### Testing power

To test statistical power, we chose three TF peak data sets of varying size (4194, 11129, and 40052 peaks) and two gene sets of vaiying size (42 and 471 genes) as our base scenarios. After assigning the peaks to genes, we randomized the genes in bins of locus length to remove all true gene set enrichment signal while keeping locus length association, and then randomly added peaks into the gene set to simulate enrichment. We chose three scenarios of enrichment, each with varying levels (*x*%=5,10,20, or 30) of enrichment:

1. CEbias: Enriched to closely satisfy the assumptions of the binary (ChIP-Enrich) model. We added peaks to *x*% of the remaining genes in the gene set without a peak. This increases the proportion of genes with a peak, without causing a large increase in the mean number of peaks per gene.

2. PEbias: Enriched to closely satisfy the assumption of the count-based (Poly-Enrich) model. We added a number of peaks, equal to *x*% of the number of peaks in the gene set, to a fraction of the genes in the gene set. This increases the mean number of peaks per gene, with little effect on the proportion of genes with a peak.

3. Balanced: We added a number of peaks, equal to *x*% of the number of peaks in the gene set, into the gene set weighted by gene locus length. This increases both the proportion of genes with a peak and the mean number of peaks per gene by a similar degree.

### Defining the true positive transcription factor-GO term pairs

For each transcription factor, we identified the gene that codes for it, and then identified every GO biological process that gene is assigned to. This set of GO terms, along with all of its ancestors, is what we use as the true positive set.

### Hybrid test

Given *n* tests that test for the same hypothesis, the same Type I error rate, and converted to p-values *p*_1_, …, *p*_*n*_, the Hybrid p-value is computed as: *P*_*hybrid*_ = *n* × min (*p*_1_, …, *p*_*n*_). This hybrid test will have at most the same Type I error rate as the *n* tests, and if at least one test is consistent (power converges to 1 as sample size reaches infinity), the hybrid test will also be consistent. Proofs and simulations of the test in general were done by Zhang et. al [13]. Here, we’ve implemented the hybrid test for users to use two methods (*n* = 2): ChIP-Enrich and Poly-Enrich. Users may also choose any two results files and run a hybrid test based on those.

### Clustering and heatmaps

For every GO term, we calculated the difference in −log_10_ p-value for each of the 90 experiments between ChIP-Enrich and Poly-Enrich, with positive values indicating a more significant result for Poly-Enrich. We then focused on GO terms where > 10% of the experiments had an absolute log_10_ p-value difference greater than 2. Clustering was performed using uncentered correlation as the similarity metric and average linkage as the clustering method. Using Java TreeView, we extracted specific groups of GO terms that contain certain strings such as “cell cycle” or “positive regul.”

### Repetitive elements

Data was obtained from the UCSC Table Browser with RepeatMasker 3.0 on the hg19 genome. We chose the two most abundant families in the dataset: Alu and L1, as well as four methods of peak-to-gene assignments: Intron, Nearest TSS, >5kb, and <5kb. Poly-Enrich was then used to perform gene set enrichment. Before clustering for the heatmap, we filtered out GO terms where there were 2 or fewer significant FDR values among the 8 categories. The clustering method was the same as mentioned in the previous section.

### Website and Bioconductor updates

The Chip-Enrich website (http://chip-enrich.med.umich.edu) updated from the *chipenrich* package version 1.7.2 to version 2.5.0. (from https://github.com/sartorlab/chipenrich, on Aug 8th, 2018). We have added the following reference genomes: human (hg38), rat (rn5, rn6), *Drosophilla melanogaster* (dm6) and zebrafish (danRer10) species. We also added the following databases from MSigDB (Version 6.0): Hallmark, Immunologic, MicroRNA, Transcription Factors and Oncogenic [6, 29]. Finally, we added sets of genes that are known to be affected by particular environmental toxins from the Comparative Toxicogenomics Database (CTD) [30].

In addition to the previous locus definitions: ‘nearest TSS’, ‘nearest gene’, ‘≤1 kb from TSS’ and ‘≤5 kb from TSS’, we also now support gene locus definitions for regions <10 kb from a TSS and gene distal regions (>10kb upstream of a TSS).

## Acknowledgements

This work was partially funded by National Institutes of Health grants R01 CA158286 and the Michigan Lifestage Environmental Exposures and Disease (M-LEEaD) Center P30 ES017885.

## Supplementary Figures and Tables

**Supplementary Figure 1:** Poly-Enrich Type I error rate plots using the <5kb and Nearest TSS gene locus definitions. Shown are the proportion of significant gene sets out of 50,150 randomized gene sets for each of 90 ENCODE ChIP-Seq dataset. Type I error rates are acceptable at the 0.05 and 0.001 level. There is some inflation for the *bylocation* randomization.

**Supplementary Figure 2:** Statistical power comparisons between ChIP-Enrich (blue) and Poly-Enrich (red) for datasets with three different sizes (i.e. number of peaks: small, medium, and large) and two gene set sizes (small and large GS), under two significance levels: α = 0.05 (A) and 0.001 (B), and three different methods of simulated enrichment (CEbias: add peaks according to the regulatory assumptions of ChIP-Enrich, PEbias: add peaks mainly according to the assumptions of Poly-Enrich, Balanced: adding peaks in proportion to each gene’s locus length). The values on the X-axis indicate the percent of extra peaks added to simulate enrichment; a higher value simulates stronger enrichment. A stricter significance level results in less power, a larger gene set results in more power, and a larger dataset (more noise) results in less power. In actuality, larger real data sets should have more power.

**Supplementary Figure 3:** Using the nearest TSS method of assigning peaks to genes, a heatmap of −log_10_ p-value differences between Poly-Enrich and ChIP-Enrich for GO terms and experiments, where each row is a GO term and each column is a ChIP-seq experiment. Red indicates Poly-Enrich was more significant, and blue indicates ChIP-Enrich was more significant. Light grey indicates the transcription factor used in the experiment was not assigned to the GO term and is omitted in the clustering. Representative GO terms are used to label each cluster.

**Supplementary Figure 4:** Enrichment results using Poly-Enrich for Alu (first four columns) and L1 (last four columns) repetitive element families using four different peak-to-gene assignments. Shown are signed −log10 FDR, where positive values (red) indicate enrichment and negative values (blue) indicate depletion. GO terms with “neuro” are almost all being significantly depleted.

**Supplementary Figure 5:** (A) A comparison of the spline estimates with and without the Gene Set (GS) covariate for a gene set with ∼1900 genes. There is very little difference despite this gene set being larger than 97% of all gene sets. (B) Signed −log10 P-value comparisons between ChIP-Enrich and its faster counterpart which usees a spline approximation. (C) Signed −log10 P-value comparisons between Poly-Enrich and its faster counterpart which uses a spline approximation. As the gene set specific spline is almost similar to the approximated spline, we see that using the spline approximation does not change the results by much, with an almost perfect identity trend with r^2^ = 1.00.

**Supplementary Figure 6:** Signed −log_10_ p-value comparison plots between ChIP-Enrich (A) and Poly-Enrich (B) vs their Score test counterparts. The gene sets that are enriched have similar significance, but those for the gene sets that are depleted may vary by several magnitudes.

**Supplementary Table 1:** Complete table of signed −log_10_ p-values for all combinations of GO terms, locus definitions (peak-to-gene assignments), and type of repeated elements. Column names are in the format of [repeated element]-[locus definition].

**Supplementary Table 2:** All 90 ENCODE datasets used for gene set enrichment testing. Downloaded from ENCODE Analysis data at UCSC: http://hgdownload.cse.ucsc.edu/goldenPath/hg19/encodeDCC/wgEncodeAwgTfbs Uniform/

