## Supplementary Figure for "Poly-Enrich: Count-based Methods for Gene Set Enrichment Testing with Genomic Regions"

$$\alpha=0.05$$

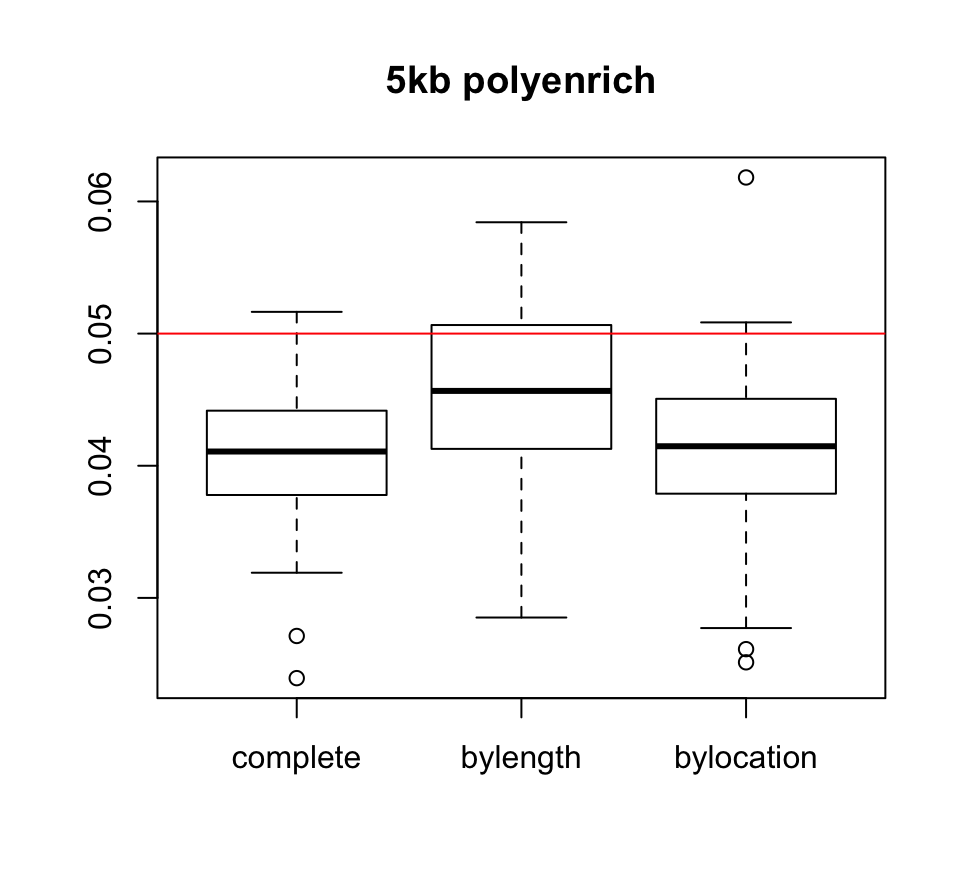

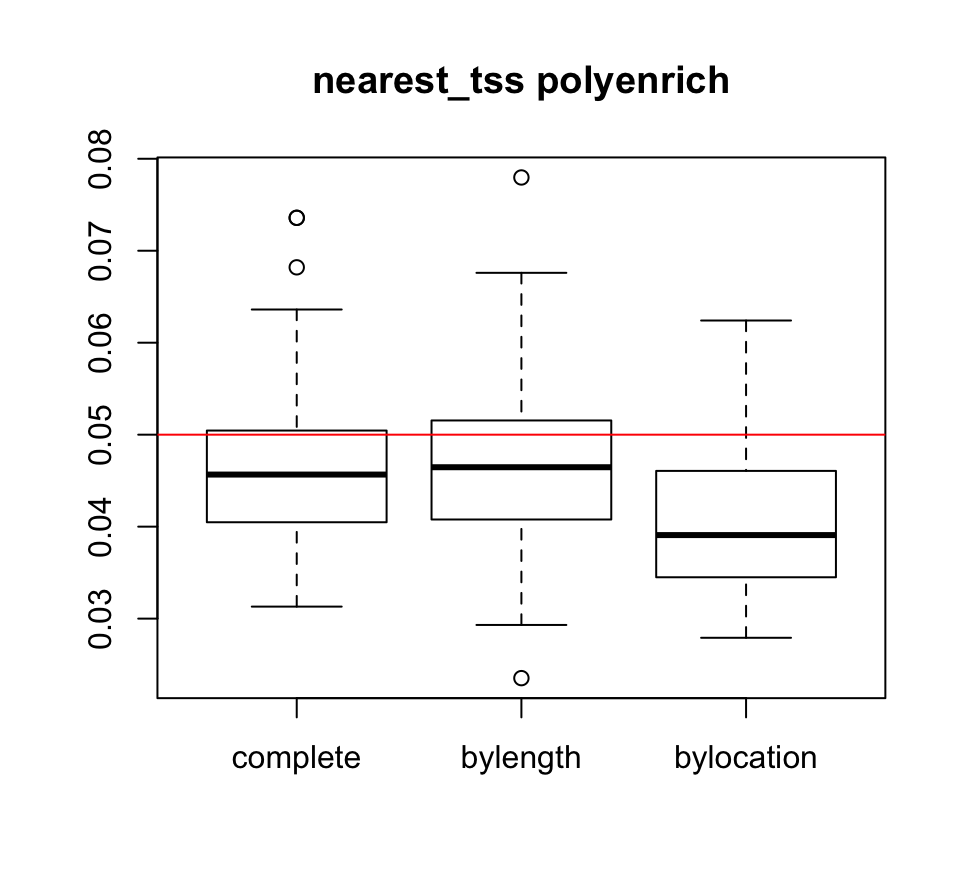

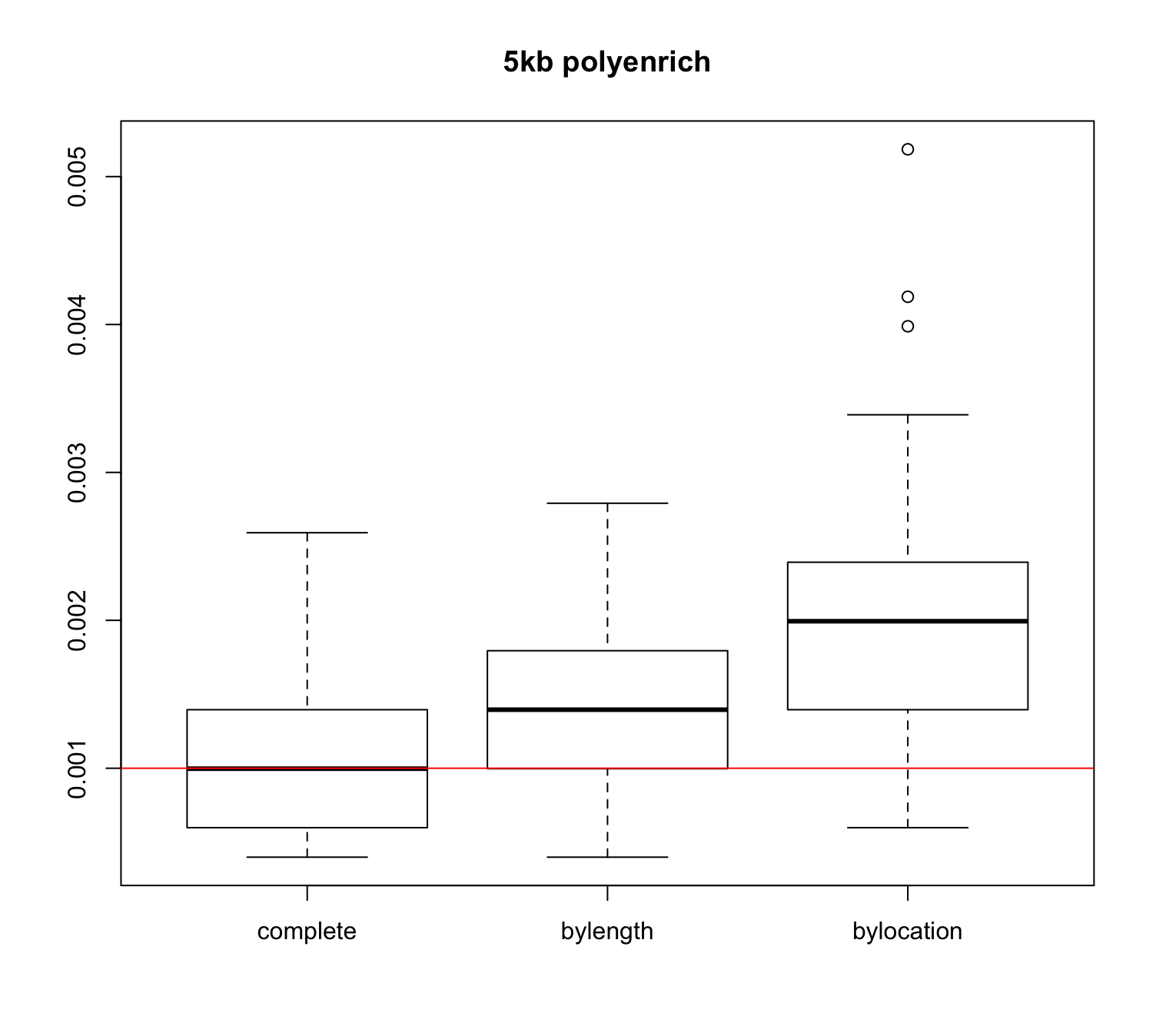

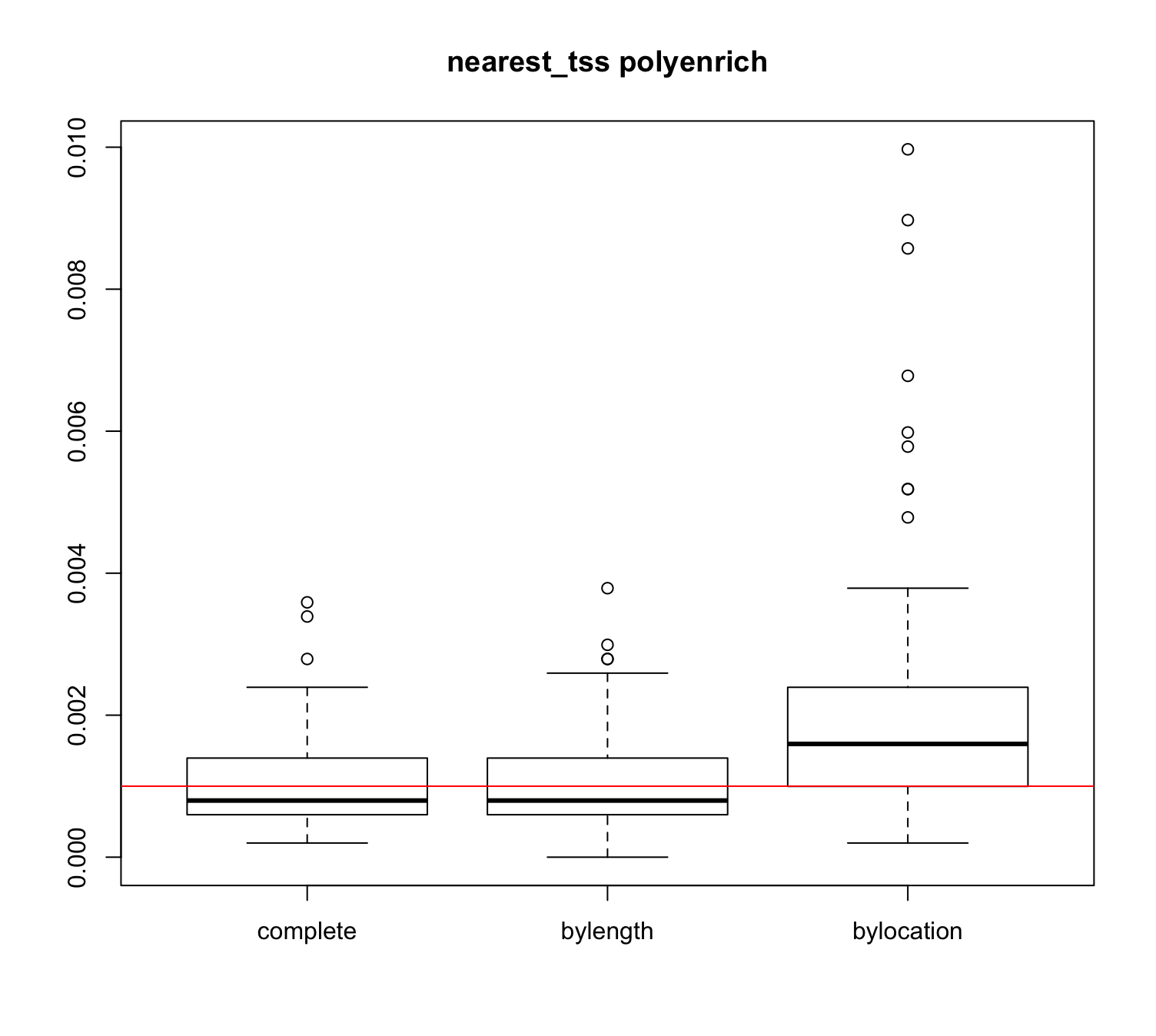

**Supplementary Figure 1:** Poly-Enrich Type I error rate plots using the <5kb and Nearest TSS gene locus definitions. Shown are the proportion of significant gene sets out of 50,150 randomized gene sets for each of 90 ENCODE ChIP-Seq dataset. Type I error rates are acceptable at the 0.05 and 0.001 level. There is some inflation for the *bylocation* randomization.

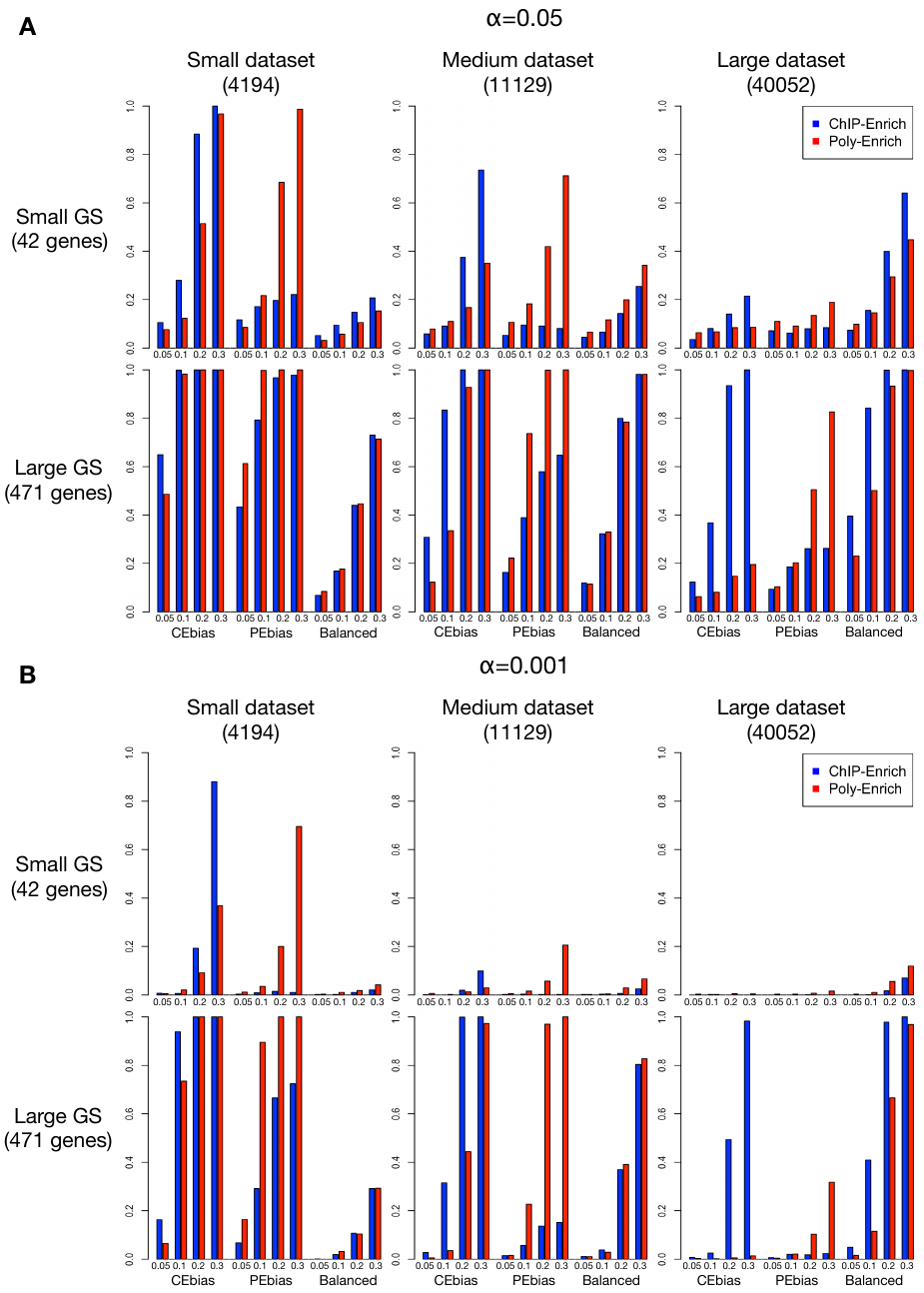

**Supplementary Figure 2:** Statistical power comparisons between ChIP-Enrich (blue) and Poly-Enrich (red) for datasets with three different sizes (i.e. number of peaks: small, medium, and large) and two gene set sizes (small and large GS), under two significance levels: α = 0.05 (A) and 0.001 (B), and three different methods of simulated enrichment (CEbias: add peaks according to the regulatory assumptions of ChIP-Enrich, PEbias: add peaks mainly according to the assumptions of Poly-Enrich, Balanced: adding peaks in proportion to each gene’s locus length). The values on the X-axis indicate the percent of extra peaks added to simulate enrichment; a higher value simulates stronger enrichment. A stricter significance level results in less power, a larger gene set results in more power, and a larger dataset (more noise) results in less power. In actuality, larger real data sets should have more power.

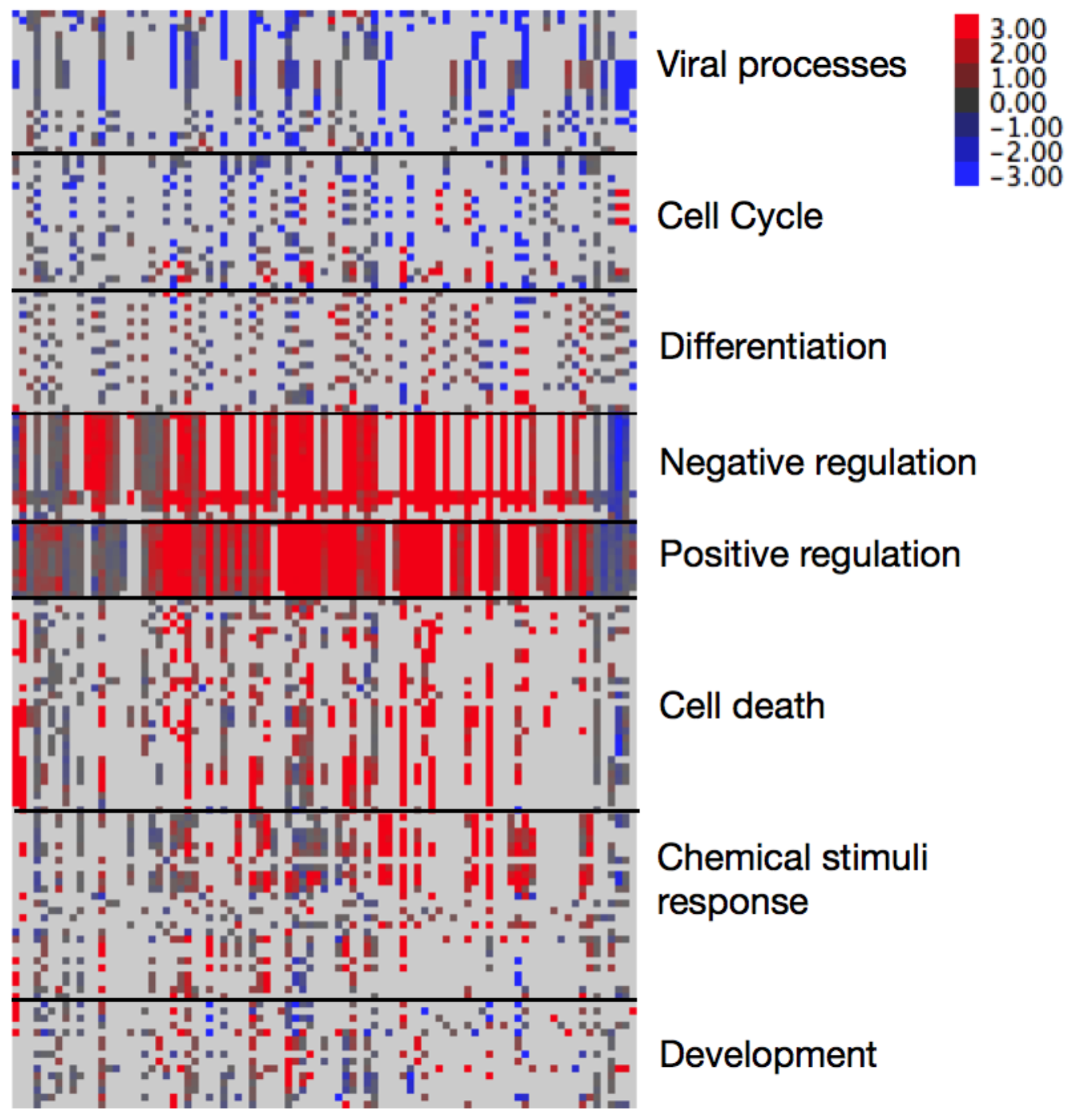

**Supplementary Figure 3:** Using the nearest TSS method of assigning peaks to genes, a heatmap of –log_10_ p-value differences between Poly-Enrich and ChIP-Enrich for GO terms and experiments, where each row is a GO term and each column is a ChIP-seq experiment. Red indicates Poly-Enrich was more significant, and blue indicates ChIP-Enrich was more significant. Light grey indicates the transcription factor used in the experiment was not assigned to the GO term and is omitted in the clustering. Representative GO terms are used to label each cluster.

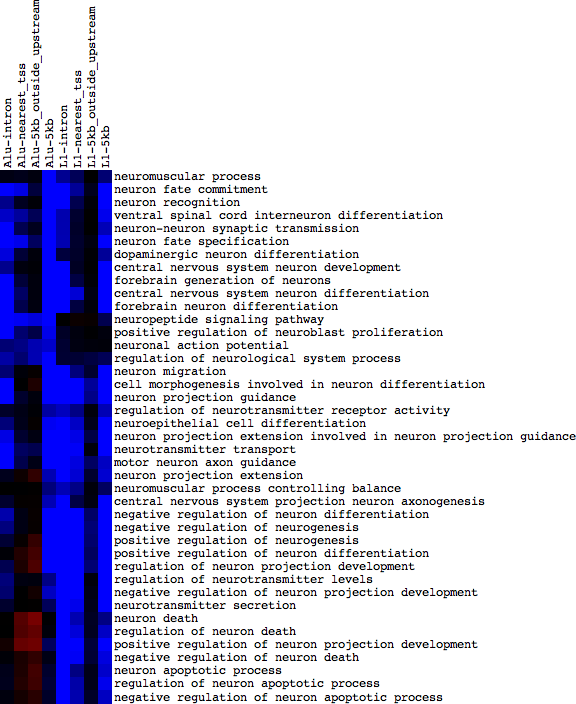

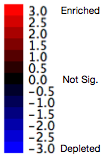

**Supplementary Figure 4:** Enrichment results using Poly-Enrich for Alu (first four columns) and L1 (last four columns) repetitive element families using four different peak-to-gene assignments. Shown are signed –log10 FDR, where positive values (red) indicate enrichment and negative values (blue) indicate depletion. GO terms with “neuro” are almost all being significantly depleted.

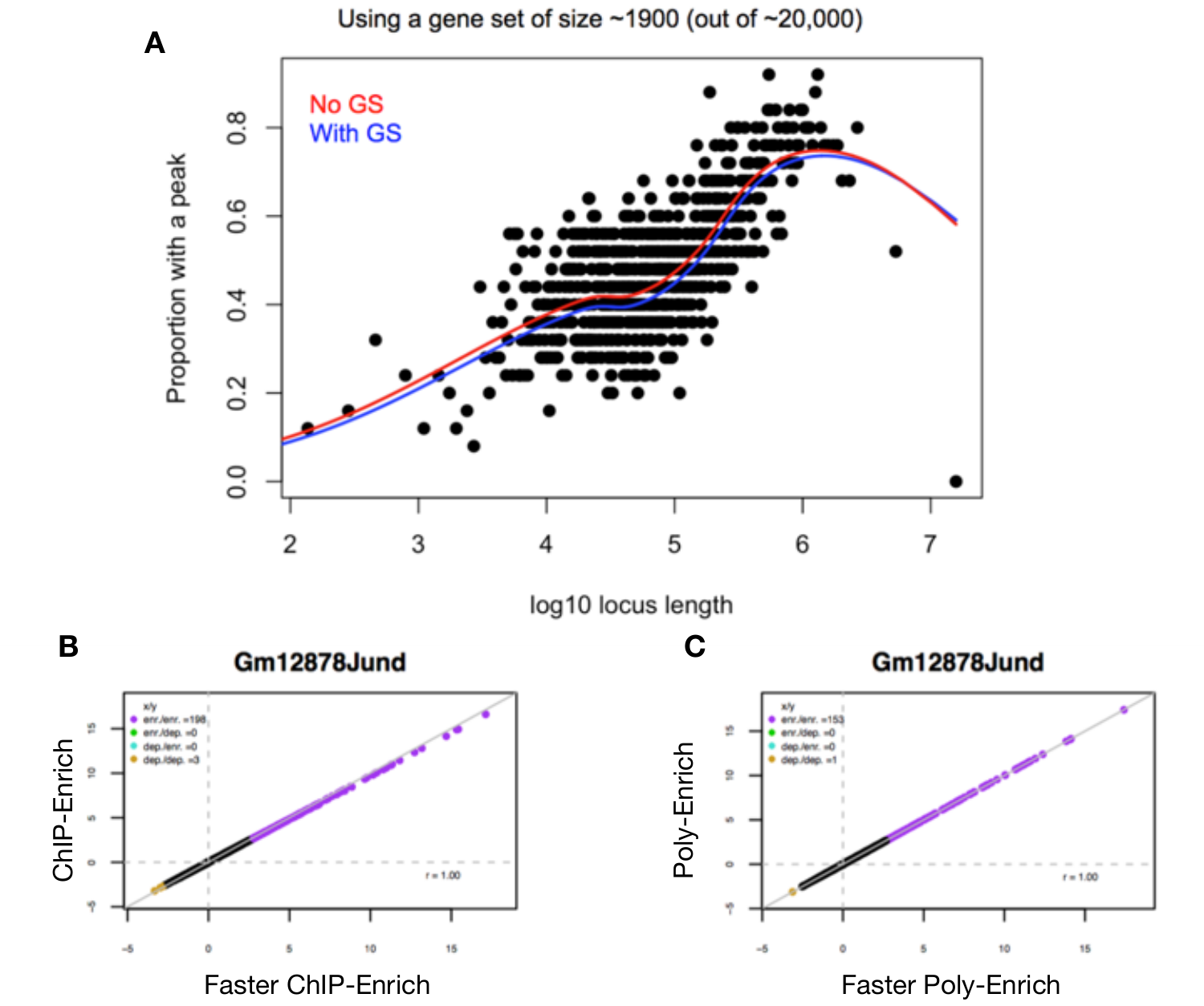

**Supplementary Figure 5:** (A) A comparison of the spline estimates with and without the Gene Set (GS) covariate for a gene set with ~1900 genes. There is very little difference despite this gene set being larger than 97% of all gene sets. (B) Signed -log10 P-value comparisons between ChIP-Enrich and its faster counterpart which usees a spline approximation. (C) Signed -log10 P-value comparisons between Poly-Enrich and its faster counterpart which uses a spline approximation. As the gene set specific spline is almost similar to the approximated spline, we see that using the spline approximation does not change the results by much, with an almost perfect identity trend with r^2^ = 1.00.

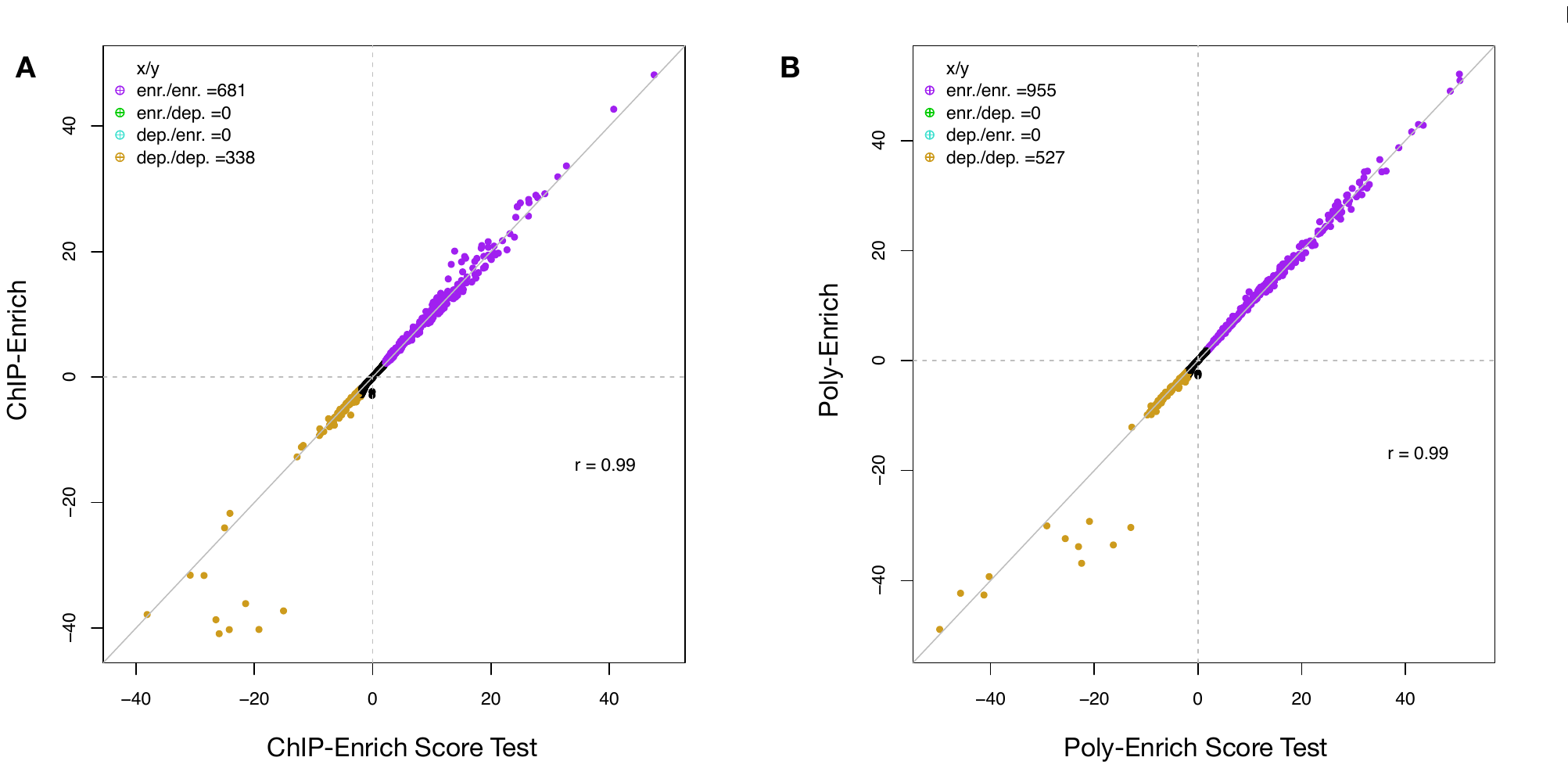

**Supplementary Figure 6:** Signed -log_10_ p-value comparison plots between ChIP-Enrich (A) and Poly-Enrich (B) vs their Score test counterparts. The gene sets that are enriched have similar significance, but those for the gene sets that are depleted may vary by several magnitudes.

**Supplementary Tables**

**Supplementary Table 1:** Complete table of signed –log_10_ p-values for all combinations of GO terms, locus definitions, and type of repeated elements. Column names are in the format of [repeated element]-[locus definition].

**Supplementary Table 2:** All 90 ENCODE datasets used for gene set enrichment testing. Downloaded from ENCODE Analysis data at UCSC: http://hgdownload.cse.ucsc.edu/goldenPath/hg19/encodeDCC/wgEncodeAwgTfbsUniform/
